# GBS-SBG - GBS Serotyping by Genome Sequencing

**DOI:** 10.1101/2021.06.16.448630

**Authors:** Suma Tiruvayipati, Tan Wen Ying, Timothy Barkham, Swaine L. Chen

## Abstract

Group B Streptococcus agalactiae (GBS; Streptococcus agalactiae) is the most common cause of neonatal meningitis and a rising cause of sepsis in adults. Recently, it has also been shown to cause foodborne disease. As with many other bacteria, the polysaccharide capsule of GBS is antigenic, enabling its use for strain serotyping. Recent advances in DNA sequencing have made sequence-based typing attractive (as has been implemented for several other bacteria, including *Escherichia coli, Klebsiella pneumoniae* species complex, *Streptococcus pyogenes*, and others). For GBS, existing WGS-based serotyping systems do not provide complete coverage of all known GBS serotypes (specifically including subtypes of serotype III), and none are simultaneously compatible with the two most common data types, raw short reads and assembled sequences. Here, we create a serotyping database (GBS-SBG, GBS Serotyping by Genome Sequencing), with associated scripts and running instructions, that can be used to call all currently described GBS serotypes, including subtypes of serotype III, using both direct short-read- and assembly-based typing. We achieved higher concordance using GBS-SBG on a previously reported data set of 790 strains. We further validated GBS-SBG on a new set of 572 strains, achieving 99.8% concordance with PCR-based molecular serotyping using either short-read- or assembly-based typing. The GBS-SBG package is publicly available and will accelerate and simplify serotyping by sequencing for GBS.

**DATA SUMMARY:**

1. The GBS-SBG package is open source and available for at Github under the MIT license (URL - https://github.com/swainechen/GBS-SBG)
2. Accession numbers of the sequencing reads and reference sequences used in the study from earlier reports have been provided within the article and the supplementary data
3. The WGS data for the 572 isolates used in the study is available at https://www.ncbi.nlm.nih.gov/bioproject/PRJNA293392

## INTRODUCTION

Group B Streptococcus (GBS, also known as *Streptococcus agalactiae*) was named for its common association with mastitis in cows (agalactiae = “no milk”). It is a common colonizer of the human gastrointestinal and urinary tract, being present in up to one third of apparently healthy individuals (1–3). Since the late 19th century, it has become increasingly associated with neonatal meningitis, today representing the most common cause (4–7). Neonatal GBS infections are classified clinically into early-onset (< 7 days of age, EOD) and late-onset (> 7 days of age, LOD) disease. In addition, GBS is an increasingly common pathogen in immunocompromised and elderly adults (8–11). Recently, GBS has been shown to cause foodborne infection associated with the consumption of raw fish in otherwise healthy adults in Singapore (and likely throughout Southeast Asia) (12–14). Accordingly, GBS is also well known to colonize and infect (often resulting in severe invasive disease) multiple other species, including fish (where it has a large impact on aquaculture) as well as other mammals, amphibians, and reptiles (15–20). As an important pathogen of humans, cows, and fish, GBS is therefore of concern for public health, economic, and zoonotic reasons.

Several decades ago, a serotyping system was established (21) based on differences in antigenicity of the polysaccharide capsule. Currently, there are 10 main serotypes (Ia, Ib, II-IX); furthermore, serotype III has been subtyped into 4 subtypes (III-1 through III-4) (22). These serotypes are encoded by the *cps* locus, which consists of both conserved (*cpsD - cpsG*) and variable (*cpsG* - *cpsK*) regions; the sequence differences, primarily in the variable region, form the basis of a commonly-used PCR-based molecular serotyping scheme (23). Epidemiological studies have made clear that different serotype (and subtype) distributions are associated with different host species, disease states, and geographical distributions (24). For example, mastitis in cows is mostly caused by serotype Ia GBS in China (25), while serotype III (subtype III-3) is most common among cows in Canada (26). Furthermore, outbreaks in infected fish are largely caused by serotype Ia, Ib, and III (the latter specifically being subtype III-4 and occurring predominantly in Southeast Asia, with recent introductions into Brazil) (14,27). Additional resolution afforded by other typing systems (multilocus sequence typing, virulence gene typing, mobile genetic elements, antibiotic resistance profiles, and whole genome sequencing) have overall confirmed these initial epidemiological observations based on serotypes (19,24,28–33). Therefore, serotyping for GBS is still a useful and important method, and continues to be particularly relevant for vaccine development (24,34). Various experimental methods have been used for serotyping of bacteria, such as enzyme immunoassay, immunoprecipitation, co-agglutination, inhibition enzyme-linked immunosorbent assay, latex agglutination, and fluorescence microscopy (21,35–39). These traditional methods have mostly been replaced by genotyping methods, often based on polymerase chain reaction (PCR)-based assays (40–43). Given recent advances in the throughput, availability, and affordability of sequencing technologies, whole genome sequencing (WGS) has now become a practical method (with advantages for automation and scale) to call serotypes in multiple bacteria, such as *Salmonella* (44,45), *Streptococcus pneumoniae* (46), *Escherichia coli* (47–49), *and Klebsiella pneumoniae* (50).

To date, three reports have explored using WGS to serotype GBS. The first used a database of 9 serotypes, focusing on the variable region of the *cps* locus (*cpsG* - *cpsK*); this database did not include serotype IX (51) and was designed for use with assembled genomes. The second database included all of the 10 main serotypes, including the full *cps* locus with both conserved and variable regions (52). This latter database was tested on raw short reads (using a mapping strategy) and assembled genomes (using a blast-based strategy); the mapping strategy was found to have higher concordance with latex agglutination-based serotyping (52). The final mapping strategy from this report does not seem to have been implemented in any publicly available software. A third WGS serotyping strategy employed partial gene sequences of the *cps* locus as a reference database for a short-read mapping strategy (53); the use of short (100-300 bp) partial gene sequences makes it ill-suited for analysis using an assembly-based strategy. Importantly, none of these databases included subtypes of serotype III, which were defined based on SNPs in a portion of the conserved region of the *cps* locus (22).

We thus aimed to devise a serotyping solution using WGS for GBS that would (i) include all existing serotypes, including subtypes of serotype III and (ii) enable accurate serotyping by both short-read-mapping and assembly-based strategies. We validated our method on the 790 strains reported in (52), achieving equivalent or higher concordance using either mapping- or assembly-based strategies. We further validated our method using an additional 572 strains, achieving 99.8% concordance with traditional PCR-based serotyping, WGS short read-based calling, and WGS assembly-based calling. Our software is called GBS-SBG and is available at https://github.com/swainechen/GBS-SBG.

## MATERIALS & METHODS

### Previously published GBS capsular locus reference sequences

Reference sequences representing the 10 serotypes (Ia – IX) were taken from (52); we refer to these as the “Kapatai database” (Figure 1, Table 1). The sequences in the Kapatai database range in length from 15,090bp to 17,514bp. Reference sequences for 9 serotypes (Ia – VIII) with lengths ranging from 4,477bp to 6,307bp were obtained from (51). We refer to these 9 sequences as the “Sheppard database”. Sequences representing two subtypes of serotype III (III-2 (AF332896) and III-3 (AF332897)) were obtained from (22).

**Figure 1.**
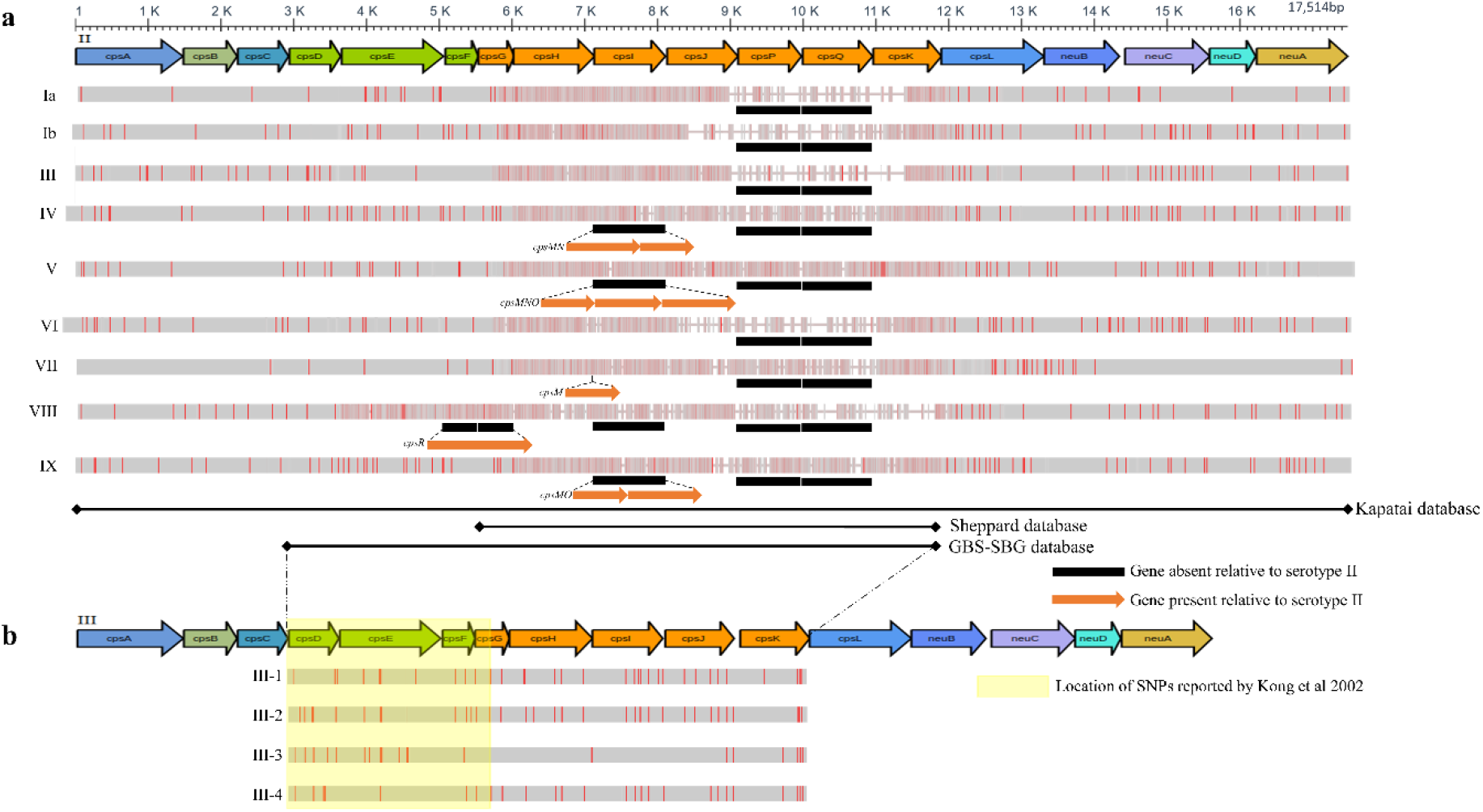
Genetic organization of the GBS cps locus and serotyping database strategies. (a) Colored arrows at the top depict the genetic organization of the serotype II sequence. Arrows are drawn to scale, as indicated by the axis at the top. Alignments for the other 9 serotype sequences (indicated by Roman numerals on the left) are shown in gray bars below. Vertical red lines indicate single nucleotide polymorphisms relative to the serotype II sequence. Black bars indicate genes that are absent from a given serotype. Orange arrows indicate genes that are unique to a given serotype, and dotted lines indicate where they fall relative to the serotype II reference. All sequences used for this figure panel are taken from (52). The black lines demarcated by diamonds at each end at the bottom of the panel indicate the region of the *cps* locus spanned by the serotype sequences in the Sheppard database, Kapatai database, and GBS-SBG databases, as indicated to the right of each line. (b) Colored arrows depict the genetic organization of the serotype III sequence from (52), using the same scale bar as for (a). Dotted lines between the panels show relative alignment of the serotype II and serotype III reference sequences. Gray bars and vertical red lines indicate alignment and SNPs for the serotype III subtypes (indicated on the left). The yellow box indicates the region in which SNPs reported by (22) are located.

**Table 1.**
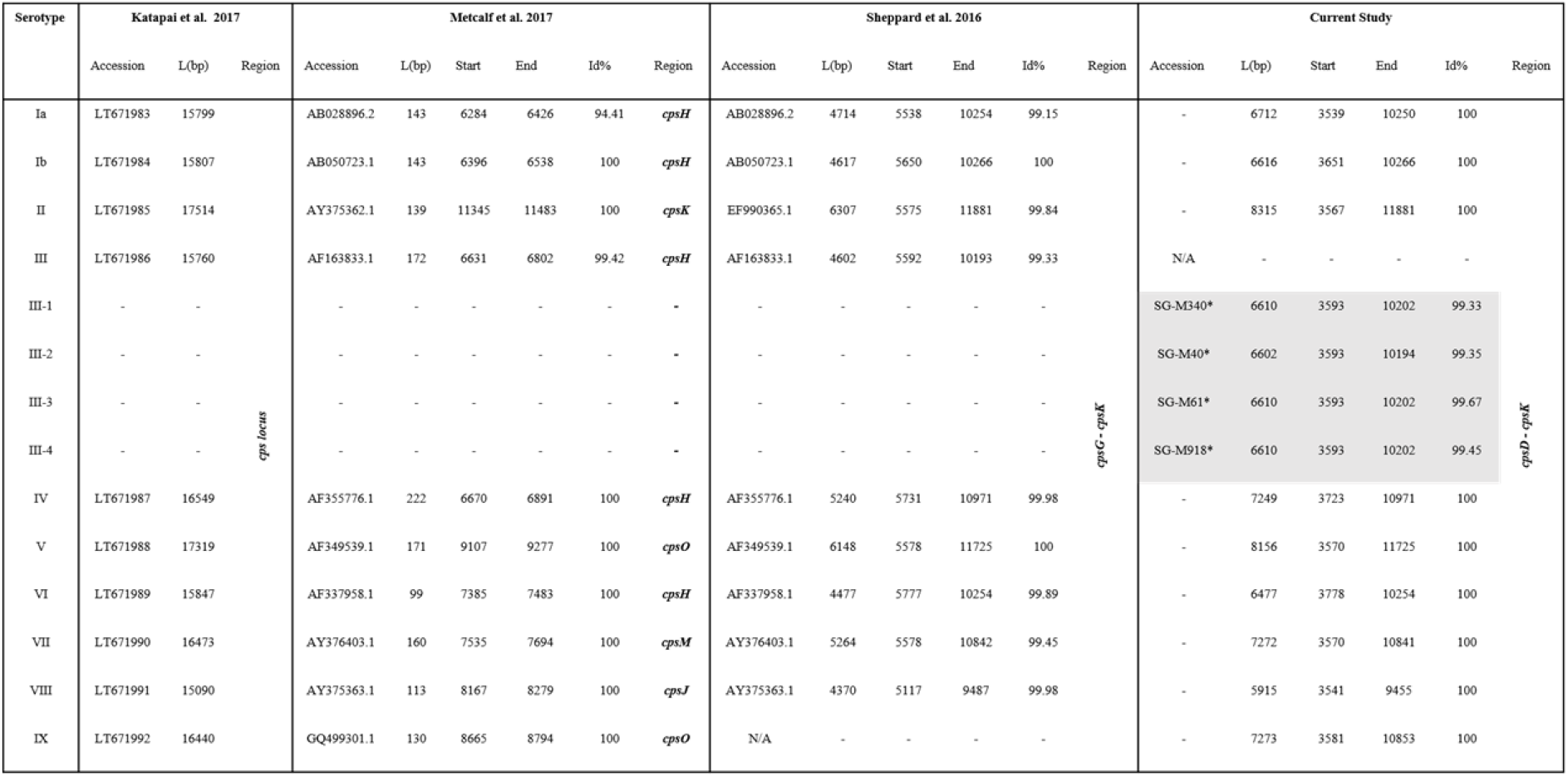
GBS serotype reference databases used in the current study. Boxes in white - Start, End, Id%, Region relative to Kapatai et al. 2017 sequences. Boxes in grey - Start, End, Id%, Region relative to serotype III Kapatai et al. 2017 sequence. (*Isolate names from the current study, Id% – Identity in percentage, L(bp) – Length in base pairs)

### GBS strains

572 GBS isolates were collected between November 2000 and July 2018. Of these, 547 were isolated from humans and 25 from fish. 487 Human isolates were collected at Tan Tock Seng Hospital (TTSH), Singapore, and stored at -70°C. All isolates were from nonpregnant adults, as TTSH does not offer paediatric or obstetric care. The study was approved by the Institutional Review Board of TTSH, Singapore, DSRB2016/00202. 45 (human), 6 (fish), 20 (1 human, 19 fish), 14 (human) isolates were obtained/ collected from Laos, Malaysia, Thailand, and Vietnam, respectively.

### PCR-based molecular serotyping

GBS isolates were subcultured onto blood agar and re-identified using a MALDI-TOF system (Bruker). GBS DNA was extracted with the EasyMag system (Biomerieux) according to the manufacturer’s instructions. Molecular serotyping of the GBS isolates was performed using 2 PCRs: “Multiplex PCR 1” and “Multiplex PCR 2” as described by Poyart *et al*. (54), with segregation into capsular polysaccharide serotypes Ia, Ib, II, III, IV, V, VI, VII, and VIII achieved using agarose gel electrophoresis based on amplicon size. If an isolate remained untypeable after these two PCRs, a third PCR step with another multiplex PCR as described by Imperi *et al*. (23) was then performed.

### Whole genome sequencing and analysis

Genomic DNA was extracted from overnight cultures on blood agar using the QIAamp DNA mini kit (cat. no. 51306). Whole-genome sequencing was performed by the Next Generation Sequencing Platform at the Genome Institute of Singapore, as previously described (13). Briefly, sequencing library preparation was done with the use of the TruSeq Nano DNA LT Library Prep Kit (Illumina) or the Nextera XT Library Prep Kit (Illumina) according to the manufacturer’s instructions. The sequencing libraries were sequenced using a NextSeq 500 or HiSeq 4000 sequencer with 2×151-bp reads (Illumina, San Diego, CA, USA). All sequencing data is uploaded in GenBank under BioProject PRJNA293392.

SRST2 version 0.2.0 (55) was used to call multilocus sequence types (MLST) using reference sequences downloaded from PubMLST (https://www.pubmlst.org). SRST2 was also used to call serotypes using a mapping strategy using the GBS-SBG database as a custom reference database, according to the SRST2 documentation. The default minimum percentage coverage cutoff (--min-coverage option; default 90%) for gene reporting was used.

Raw short read (FASTQ) sequences were also assembled with velvet (version-1.2.10) (56) using the VelvetOptimizer helper script (version 2.2.4) and a minimum contig cutoff of 500bp, scaffolded with OPERA-LG (version 2.0.6) (57), and finished with FinIS (version 0.3) (58). Alignment-based calling of serotypes was done using a single script that performed alignment against the GBS-SBG database using BLASTN (59) and processed the results (available at https://github.com/swainechen/GBS-SBG). For consistency with SRST2, a minimum nucleotide identity of 90% across 90% of the reference serotype length (i.e. 90% coverage) was required for making a serotype call.

### Creation of the GBS-SBG database

For all serotypes except for serotype III, the sequences reported by (52) (i.e. the Kapatai database) were used (Table 1). Multiple sequence alignments were made using MAFFT version 7.45 (60). Sequence alignments were visualized using Jalview 2.11.1.3 (61). Sequences were trimmed to a uniform 5’ and 3’ end (see results) (Figure 1). For the subtypes of serotype III, reference sequences for III-2 and III-3 were taken from (22). SNPs specific for III-1 and III-4 (22) were introduced into the reference serotype III sequence from (52) (of note, this serotype III sequence (LT671986) appeared to be a hybrid between III-2 and III-4 (based on characteristic SNPs for these subtypes) and was therefore not clearly subtypeable itself (Supplementary Table 1)). To generate reference sequences for III-1 and III-4, all of the new strains that were typed as serotype III by PCR (173/572 strains) were considered. All of these were unambiguously typable as one of the four subtypes of serotype III based on the SNPs reported by (22) (Supplementary Table 1, Supplementary Figure 1). For each subtype, one corresponding assembled sequence (Table 1) was then included as a reference sequence in the final database.

### Pairwise distance and phylogenetic analysis

Pairwise distances and maximum likelihood phylogenetic analysis using a Tamura-Nei model (62) was performed using MEGA X (63). The initial tree(s) for the heuristic search were obtained automatically by applying the Neighbor-Join and BioNJ algorithms to a matrix of pairwise distances estimated using the Maximum Composite Likelihood (MCL).

## RESULTS

### Existing GBS serotyping strategies do not include all known serotypes and only work with a single data type

A valuable dataset, comprising 790 isolates with WGS data and experimental serotyping by latex agglutination, was made publicly available in the report of one of the short-read mapping GBS serotypers (52). This study achieved a concordance of 725/790 between WGS (using short read mapping) and latex agglutination. The authors also tested an assembly-based method using their same sequence database, but found lower concordance (664/790) (Supplementary Data 2). The lower concordance using assemblies, of course, meant that the concordance between a mapping- and assembly-based strategy was also not perfect (721/790).

We performed serotyping of these 790 isolates using a short read mapping strategy (with the popular SRST2 program) against the Kapatai database; this resulted in 431/790 correct serotypes. The 358 discrepant calls were all miscalls between serotypes III and Ia (355 were miscalled as Ia while 3 were miscalled as III). Of note, high similarity between the *cps* loci for serotypes Ia and III has previously been reported (52,64).

We further leveraged this data set of 790 strains to evaluate the suitability of the reference database reported in (51) (designed for an assembly-based strategy) using a mapping-based strategy. In agreement with (65), we found that this database performed very well, resulting in 780/790 correct serotype calls. Of the 10 discordant calls, 9 were serotype IX (which was not present in the Sheppard database) while 1 was miscalled as serotype III (instead of Ia).

The report by (53) was also originally designed for mapping-based typing. We did not assess its performance for assembled sequences because the reference sequences were very short (100-300 bp) (Table 1).

Therefore, none of the three existing databases was definitively usable as-is for both mapping- and assembly-based typing. Furthermore, none included the possibility of calling serotype III subtypes.

### Construction of a complete reference serotype database usable for both assembly- and short read-based analyses

We therefore sought to construct a single database that could provide high accuracy using both short reads and assemblies as well as provide serotype III subtyping. Previous studies have noted that the variable region of the *cps* locus is important for assembly-based typing; this was the main region used for the assembly-based method (51). Short read mapping-based typing, such as that performed by SRST2 (which was used by (53)), is typically designed to differentiate between closely related alleles (i.e. conserved regions), making it well suited for multilocus sequence typing (MLST), for which the 5’ and 3’ ends of the reference sequences are strictly trimmed to provide uniform-length alignments of the typing alleles. Interestingly, we found that using SRST2 also worked well with the Sheppard database but less well with the Kapatai database, despite the latter also including the variable region of the *cps* locus.

We therefore hypothesized that a new database should include both conserved and variable portions of the *cps* locus, in order to accommodate both short read- and assembly-based typing. We also sought to align the start and end of each sequence. We built upon the database created by (52), as it already achieved good performance with short read mapping (though not with SRST2).

We constructed a new database as follows (see Methods for more details):

1. Take all 10 main serotypes from the Kapatai database and align them.
2. Trim all sequences on the 5’ end to the region that includes the SNPs reported to differentiate serotype III subtypes (22).
3. Trim all sequences on the 3’ end to the edge of the variable sequence, which corresponds to the right edge of the sequences in the Sheppard database; this resulted in a ∼8kbp reference sequence for each serotype.
4. Replace the single serotype III sequence with representatives from the four subtypes (see below).

The serotype III subtypes have been defined by SNPs that are found in a conserved region of the *cps* locus (22). Full length sequences for these have not been explicitly identified, however. We therefore used the WGS assemblies for our newly sequenced strains (see below) that were typed as serotype III by PCR (173 strains in total). These were unambiguously classifiable into the four subtypes based on the SNPs reported by (22), resulting in 26 III-1, 22 III-2, 8 III-3, and 117 III-4 isolates. For each subtype, we then took the consensus sequence over the ∼8kbp typing region (for each subtype, this consensus sequence exactly matched the sequence of multiple assemblies in our data set) and included this in our new reference serotyping database. We refer to this as the GBS-SBG database.

### The GBS-SBG database enables serotyping by both mapping and assembly-based strategies

We tested the GBS-SBG database on the 790 isolates from (52), using both short read- (with SRST2) and assembly-based (using BLASTN, with 90% identity and 90% coverage cutoffs; see Methods) strategies. We used the mapping-based calls from Supplementary Data 2 in the original report (52) as the gold standard (as noted above, these mapping-based calls were only concordant with assembly-based calls for 721/790 strains in the original report). We found 100% concordance (790/790) and 94.6% (748/790) with the reported serotypes when using read mapping (SRST2) and assembled sequences (BLASTN), respectively, with the GBS-SBG database. The differences in the 42 strains from the assembly-based analysis were all due to calls as non-typeable. Further examination showed that these 42 strains had lower sequencing depth (p < 1.044e-12, 2-tailed Mann-Whitney U-test; Supplementary Figure 4), which led to an inability to assemble portions of the *cps* locus. The regions that were not assembled correlated with areas of low coverage, which still largely remained above the minimum read depth (5x) required by SRST2 to make a call (Supplementary Figure 5). Use of the original assemblies by (52), which were performed using SPAdes instead of velvet, led to a different set of nontypeable strains, which also had generally lower coverage (Supplementary Data 2).

### Validation of GBS-SBG on an a previously unanalyzed set of strains

A total of 572 previously unanalyzed GBS isolates were serotyped by PCR and whole genome sequenced. By PCR, none of these were typed as serotype VIII, and one isolate (SG-M122) was non-typeable (Supplementary Data 3). We again used both short read-(with SRST2) and assembly-based (using BLASTN) strategies with the Sheppard and Kapatai databases as well as with our new database. The concordance with the PCR-based serotyping is summarized in Supplementary Data 3. Using the GBS-SBG database achieved the best concordance by both mapping- and assembly-based strategies (571/572, 99.8% concordance). Furthermore, we achieved 100% accuracy in calling serotype III subtypes. For assembly-based analysis only, the Sheppard database was equivalently good, with the exception of not including a serotype IX reference sequence.

Interestingly, using a mapping-based strategy (although with SRST2), the Kapatai database led to concordant calls with PCR serotyping in only 386/572 isolates. The majority of the discrepancies (168/186) were due to a miscall of serotype Ia (by WGS) for serotype III (by PCR). The remaining 18 isolates had the opposite problem (called serotype III by WGS using the Kapatai database and serotype Ia by PCR) (Supplementary Data 3). As noted above, this is consistent with the close similarity between serotype Ia and III (52,64), which we saw with the reference sequences we used as well (Figure 2, Supplementary Figure 2). Furthermore, the serotype III sequence included in the Kapatai database itself was not clearly one of the previously described subtypes, but instead appeared to be a hybrid of III-2 and III-4 (Supplementary Data 1). Regardless, these miscalls between serotypes Ia and III were resolved by using our new database, possibly in part by the inclusion of all four subtypes and by the alignment of the 5’- and 3’-ends of the reference sequence, facilitating discrimination by SRST2.

**Figure 2.**
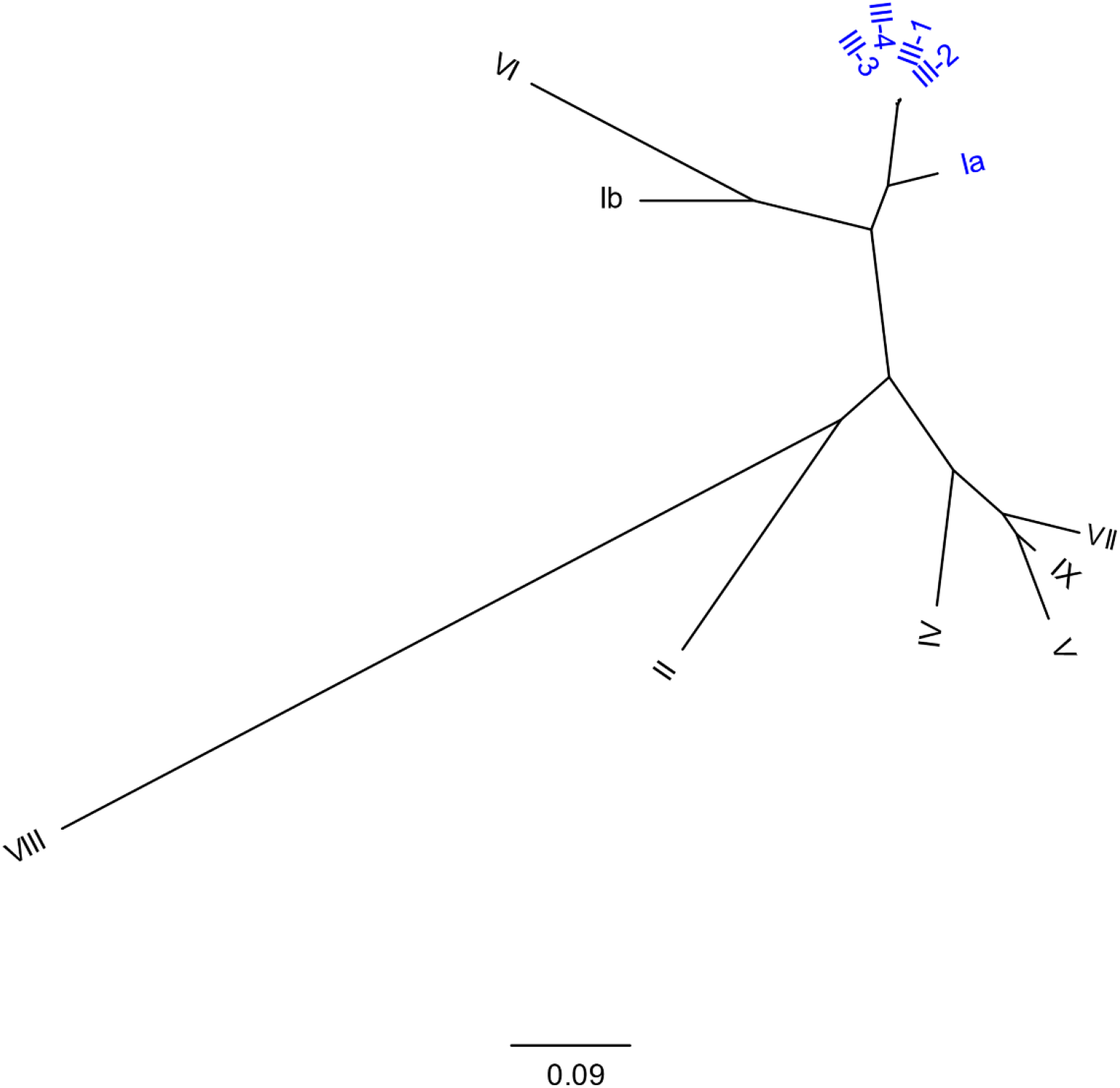
Phylogenetic relationship of serotype reference sequences in the GBS-SBG database. A maximum likelihood tree is shown; a total of 13 serotype sequences were used to plot an unrooted binary tree. The scale bar for branch length at the bottom indicates the number of substitutions per site. There were a total of 11911 positions in the alignment. The labels for serotypes Ia and subtypes of serotype III are highlighted in blue for ease of visualization.

We found only one strain (SG-M666) that had a discordant call with PCR (Ia) and WGS (V) typing (both short reads and assemblies gave a ser
otype V call). On repeat PCR typing, this strain was again called a serotype Ia. Examination of the WGS data showed that the *cps* locus was 100% identical to the serotype V reference sequence, with a 614 bp deletion affecting the *cpsN* and *cpsO* genes. As this deletion is <10% of the length of the serotype V reference sequence, GBS-SBG called this as serotype V (Supplementary Figure 3). In contrast, the SG-M666 *cps* locus was only 98.8% identical to the serotype Ia reference sequence in regions where it was aligned by BLASTN. Furthermore, it had a 3,423 bp deletion (>10% of the total length) relative to the serotype Ia reference sequence (such that the *cpsG-cpsJ* genes were missing). Closer examination of the target regions for the typing PCRs showed that the 614 bp deletion eliminated one of the priming sites for the serotype V PCR; in this multiplex PCR, serotype Ia and serotype V have two PCR products of identical size, but serotype V should have a third PCR product that overlappedthis deletion. This led to a miscall of this strain as serotype Ia by PCR (Supplementary Figure 3). Therefore, while discordant between PCR and WGS, we believe that in this case the WGS typing (serotype V) is the correct call, though the 614 bp deletion may mean this strain is actually nontypeable by latex agglutination.

## DISCUSSION

The dramatic increase in availability of whole genome sequencing has increased the importance of *in silico* approaches for analysis of multiple bacteria. Serotyping remains an important epidemiological adjunct for many bacteria of public health importance. For GBS, three previous studies have reported WGS-based serotyping, two using a mapping approach (one using SRST2 and another using a custom SNP-based method) (52,53) and one using an assembly approach (51). Having a database that is compatible with a general tool like SRST2 (as opposed to a custom mapping method) provides advantages for enabling standard analysis of multiple bacteria, such as for MLST, antibiotic resistance, and virulence genes. SRST2 also includes a reference database enabling serotyping of *E. coli*. However, many WGS-based bacterial serotyping methods (such as for *E. coli, K. pneumoniae*, Salmonella, Group A *Streptococcus*, etc.) still use an assembly-based analysis (50,65–67).

Furthermore, long-read sequencing technologies are also becoming more popular, which likely will be better suited to assembly-based methods. We therefore speculate that both short read- and assembly-based methods will be important for some time; however, finding a common reference database which works equivalently well for mapping- and assembly-based serotype calling for GBS has been challenging (52).

Here, we have developed a single database (GBS-SBG) that provides the highest accuracy (as assessed by concordance with PCR-based molecular serotyping) for serotyping GBS regardless of whether a mapping- or assembly-based strategy is used. This database can be used directly by the popular SRST2 program. Previous studies had already noted that use of the full *cps* locus, the conserved *cpsD-G* genes (51), or the variable *cpsG-K* genes (52,53) were not simultaneously suitable for use with both mapping- and assembly-based analyses. Therefore, we included portions of both the variable and conserved regions. Inclusion of a part of the conserved region enabled us to further incorporate the discriminating SNPs for subtypes of serotype III (22). Finally, to match other databases provided with SRST2, we ensured that the reference sequences had aligned 5’- and 3’-edges.

We validated the accuracy of serotyping with this database using a previously published dataset of 790 sequenced strains (typed with latex agglutination) (52), achieving equal or higher concordance than the original report, regardless of whether a mapping- or assembly-based approach was used. Calls based on analysis of assemblies appear more sensitive to sequencing depth, with lower sequencing depth correlating with an increase in non-typeable calls. To handle these cases, GBS-SBG can be set to report the next best call (i.e. with a coverage < 90%), which were all concordant with the mapping-based analysis and the gold standard serotypes reported by (52). We further validated the accuracy with a new dataset of 572 sequenced strains (typed by PCR), achieving 99.8% (571/572) concordance using either analysis method. On careful examination, the single discordance appeared to be caused by an error in the PCR-based serotyping, due to a 614 bp deletion in an otherwise canonical serotype V locus that affected the priming sites for one of the PCRs. It remains possible that the WGS call (serotype V) is also incorrect, as the 614 bp deletion affects genes important for capsule assembly, and this strain may actually be nontypable by latex agglutination. However, this example highlights the advantage of WGS in providing more data for serotyping and in being less susceptible to sequence variants or mutations that affect PCR priming sites.

We attempted to use data sets reported by the other two reported WGS serotyping methods (51,53). Unfortunately, serotype data for individual strains was not reported in (53); however, the number of strains for each serotype was reported in aggregate, and WGS-based serotyping using our new database, with either analysis method, reproduced a similar serotype distribution (data not shown). We were unable to locate sequencing data for the strains analyzed by (51) in public repositories.

In conclusion, we found that use of our new database enables accurate WGS-based serotyping of GBS by both mapping- and assembly-based strategies. The mapping-based strategy relies on the commonly used SRST2 tool. The ability to also use an assembly-based strategy with the same reference database increases the generality and anticipates a continued transition to long-read sequencing. Finally, this new database provides a further advantage over all previously reported databases in allowing subtyping of serotype III.

## Author contributions

Conceptualization, Methodology: S.L.C.; Software: S.T., S.L.C.; Validation: S.T., T.W.Y.; Formal Analysis: S.T., S.L.C., Investigation: S.T., T.W.Y.; Resources: T.B., S.L.C.; Data Curation: S.T., T.W.Y., T.B., S.L.C.; Writing – Original Draft Preparation: S.T., S.L.C., Writing – Review and Editing: S.T., T.B., S.L.C., Visualisation: S.T., S.L.C., Supervision: T.B., S.L.C., Project Administration: T.B., S.L.C., Funding: T.B., S.L.C.

## Acknowledgements

We would like to thank the members of the Chen lab for helpful comments and inputs. We thank Kurosh Mehershahi and the Genome Institute of Singapore’s Next Generation Sequencing Platform for assisting with library preparation and sequencing.

## Supplementary Figures

**Supplementary Figure 1.**
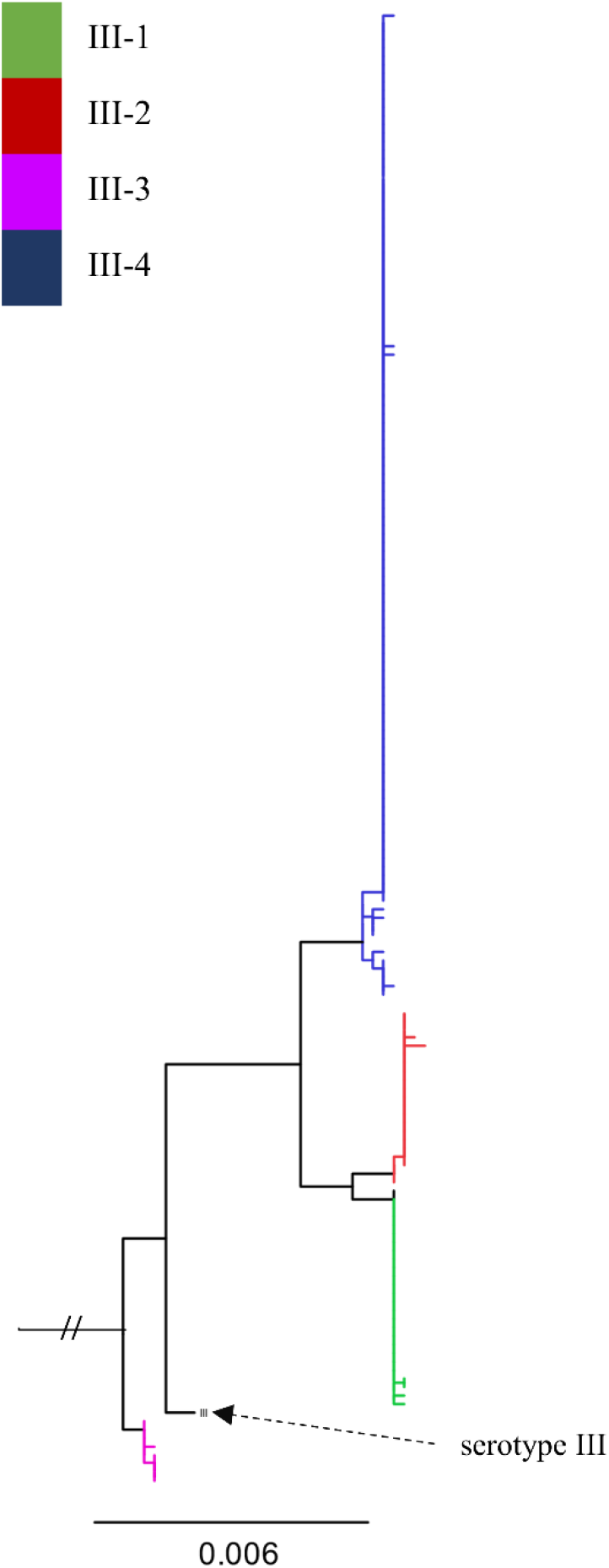
Phylogenetic tree of serotype III sequences. A maximum likelihood tree is shown; a serotype Ia sequence was used as an outgroup to root the tree. This analysis involved 175 nucleotide sequences. The scale bar for branch length at the bottom indicates the number of substitutions per site. There were a total of 6789 positions in the final dataset. Branch colors indicate the subtype of serotype III, as indicated by the legend at the top left. The sequence labelled in black and highlighted with an arrow is the reference serotype III sequence from the Kapatai database.

**Supplementary Figure 2.**
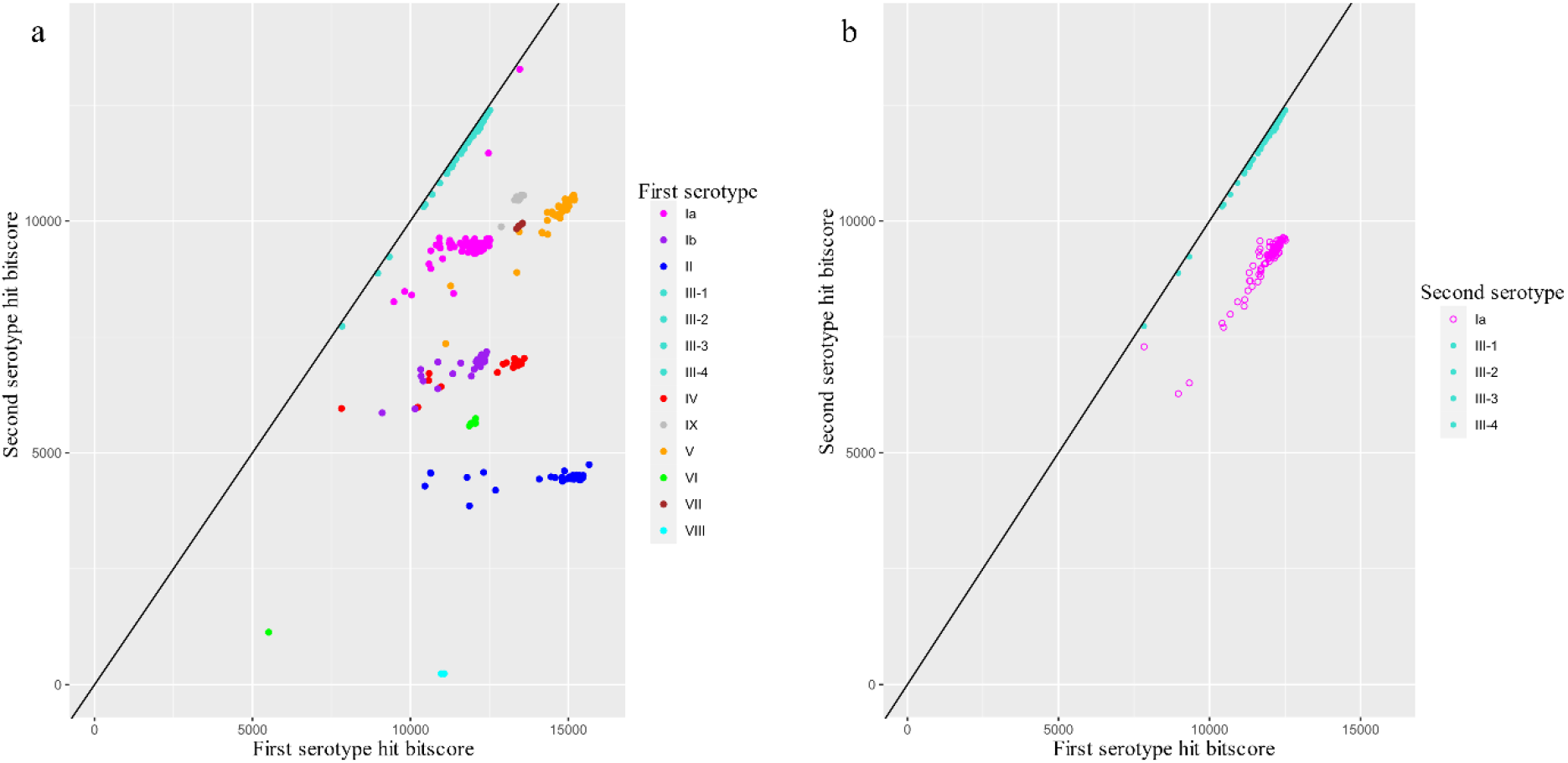
Typeability based on best versus second-best serotype match. Each graph plots the total Bitscore (reported by BLASTN) for the second best-scoring serotype (y-axis) against the total Bitscore for the best-scoring serotype matched (x-axis). A solid black line is drawn at y=x; thus, points further to the right of the black line indicate better typeability (i.e. a larger difference in total Bitscore between the best and second-best match). (a) Each data point represents one strain (n = 790 total). Points are colored based on the best serotype matched, as indicated by the legend on the right. (b) Only strains with a best match to one of the serotype III subtypes are plotted (n = 356 total). Each strain is represented by two data points: one point is as plotted in panel (a), while a second data point is plotted in which the second-best match was taken to be the best non-serotype III match. Thus, the two data points for each strain are aligned vertically. Points are colored based on the serotype of the second-best match as indicated by the legend on the right.

**Supplementary Figure 3.**
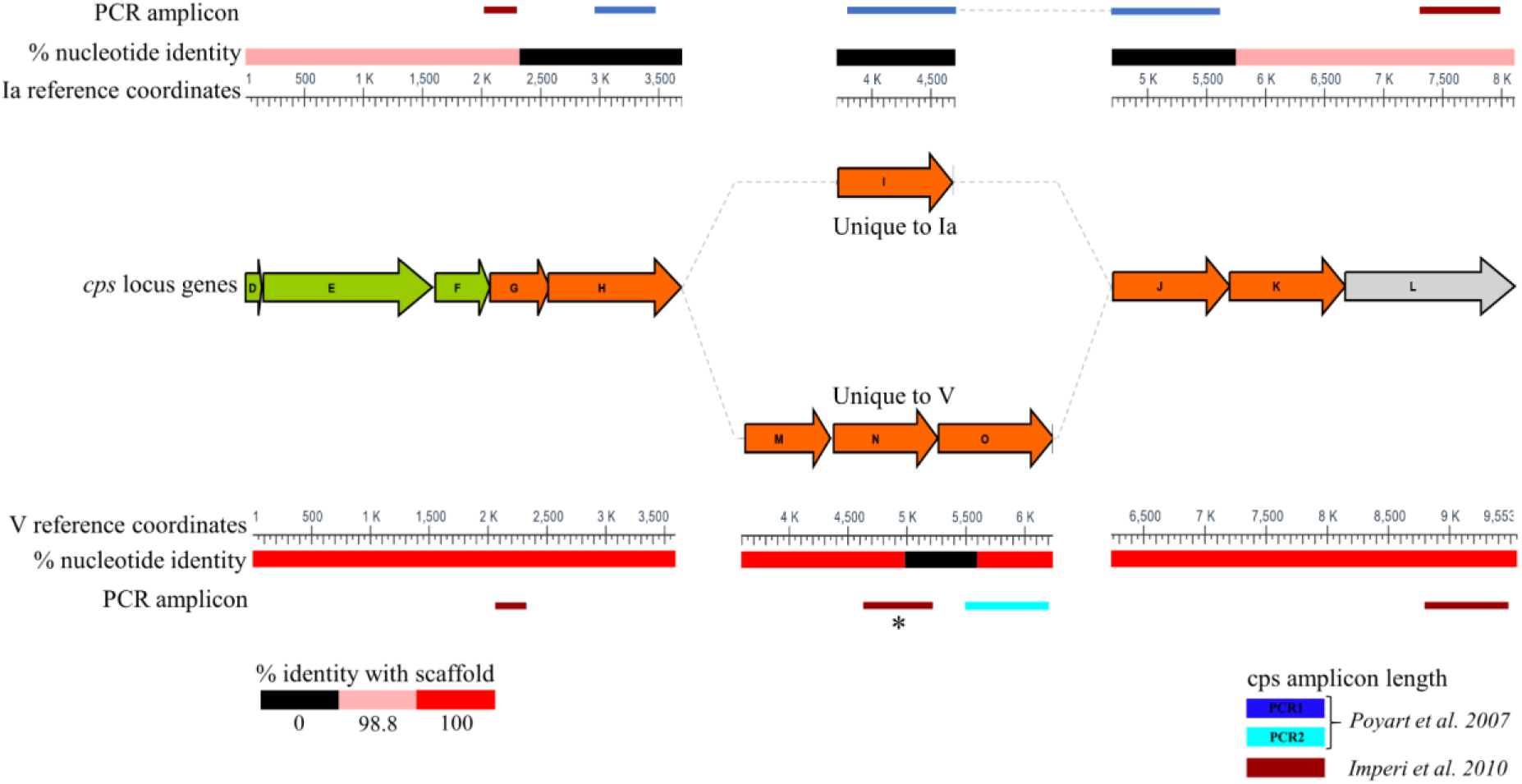
Detailed analysis of PCR versus WGS serotyping of SG-M666. Colored arrows in the middle track depict the genetic organization of the reference sequences for serotype Ia and V. Colors for each gene are the same as shown in Figure 1. Genes conserved between serotype Ia and V are shown in the middle. Genes unique to serotype Ia and serotype V are shown offset above and below, respectively, the conserved genes. Dotted lines join contiguous sequences, which are offset due to different lengths of unique genes. A nucleotide scale bar for each serotype is shown (Ia above, V below). Solid colored bars adjacent to the nucleotide scale indicate the percent nucleotide identity (as reported by BLASTN) between SG-M666 and the serotype Ia and serotype V reference sequences, according to the color legend at the bottom left. The location of the PCR amplicons used for PCR-based serotyping (Poyart C, et al. 2007, Imperi M, et al. 2010) are shown as colored lines above (for serotype Ia) and below (for serotype V) the nucleotide identity tracks; those with coordinates overlapping black bars on the “% nucleotide identity” track are predicted to yield no amplification. Of the three PCRs from Imperi, et al. 2010, the PCR that distinguishes serotype V from serotype I (at ∼4.6-5.2kb on the serotype V scale; highlighted with an asterisk) is predicted to be negative, thus mimicking a serotype I result.

**Supplementary Figure 4.**
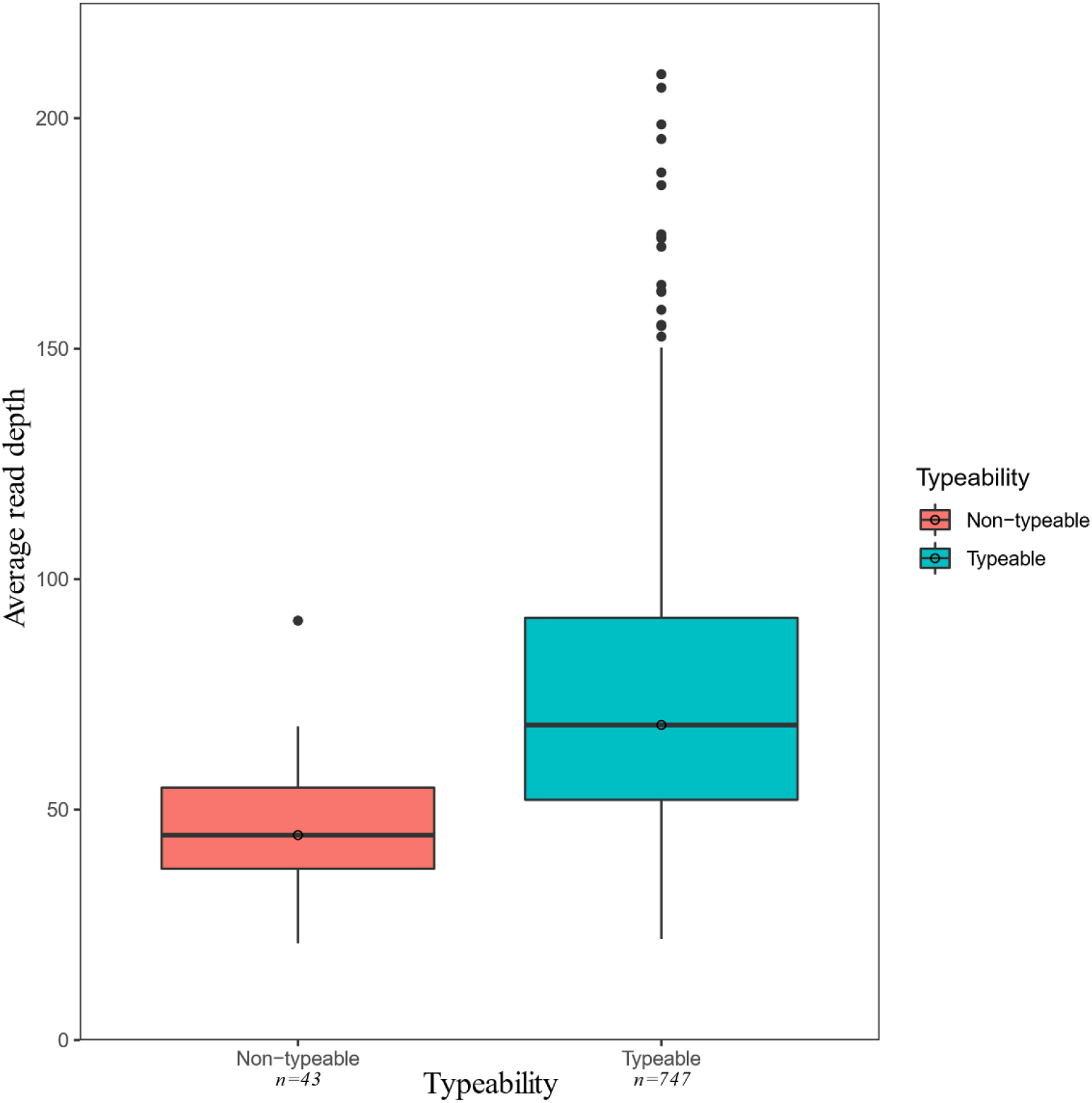
Box plot showing the distribution of the average read depths in the Kapatai datasets. The median is indicated by the horizontal line in the middle of each box and the 75th and 25th percentiles indicated by the top and bottom borders of the box, respectively. Whiskers indicate 1.5x the inter-quartile range. Outliers are represented by individual data points. The Mann–Whitney U test was used to compare the median read depths of the non-typeable (*n=43*) and typeable (*n=747*) datasets (p< 1.044e-16).

**Supplementary Figure 5.**
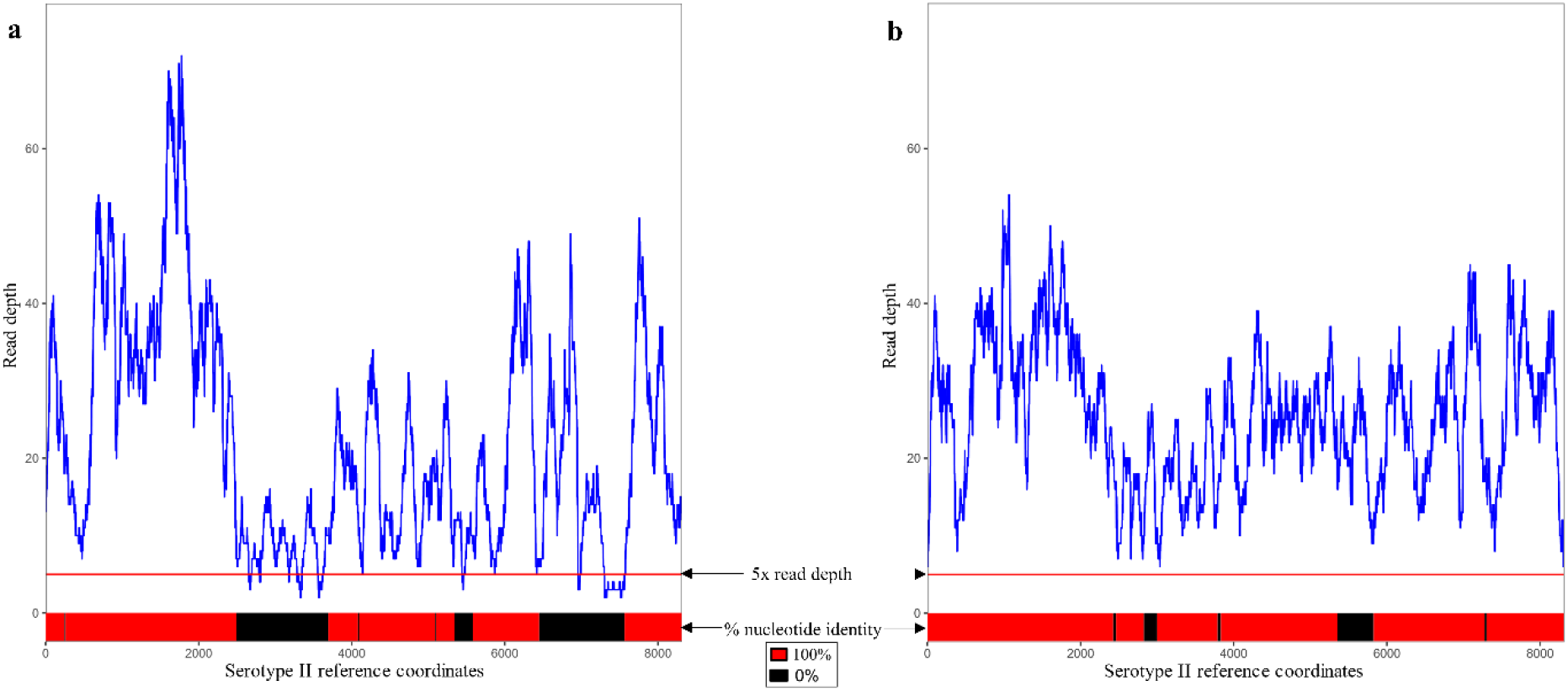
Detailed analysis of a GBS-SBG non-typeable and typeable dataset. Each graph plots the read depth per reference base (y-axis) against the serotype II reference coordinate (x-axis). A solid red line is drawn at y=5, which is the minimum read depth required by SRST2 to consider a base “covered”. (a) PHEGBS0096 (ERR1741774), which was typed as serotype II by SRST2 using short reads (20.967x average read depth). This was called as nontypeable by GBS-SBG using the assembly; the next best hit was serotype II, with 99.9% identity over 68.5% coverage. (b) PHEGBS0074 (ERR1742026) was typed as serotype II by both SRST2 (for short reads) (23.678x average read depth) and GBS-SBG (for assemblies). Note that the coverage does not fall as low as it does for PHEGBS0096, and the assembly of the cps locus is more complete. Solid colored bars at the bottom of each graph indicate the percent identity (as reported by BLASTN) between the serotype II reference sequence (8315bp) and PHEGBS0096 and PHEGBS0074 samples, according to the legend between the panels.

